# Mechanics of multi-centrosomal clustering in bipolar mitotic spindles

**DOI:** 10.1101/2019.12.17.879817

**Authors:** S Chatterjee, A Sarkar, J Zhu, A Khodjakov, A Mogilner, R Paul

## Abstract

To segregate chromosomes in mitosis, cells assemble mitotic spindle, a molecular machine with centrosomes at two opposing cell poles and chromosomes at the equator. Microtubules and molecular motors connect the poles to kinetochores, specialized protein assemblies on the centromere regions of the chromosomes. Bipolarity of the spindle is crucial for the proper cell division, and two centrosomes in animal cells naturally become two spindle poles. Cancer cells are often multi-centrosomal, yet they are able to assemble bipolar spindles by clustering centrosomes into two spindle poles. Mechanisms of this clustering are debated. In this study, we computationally screen effective forces between a) centrosomes, b) centrosomes and kinetochores, c) centrosomes and chromosome arms, d) centrosomes and cell cortex, to understand mechanics that determines three-dimensional spindle architecture. To do this, we use stochastic Monte Carlo search for stable mechanical equilibria in effective energy landscape of the spindle. We find that the following conditions have to be met to robustly assemble the bipolar spindle in a multi-centrosomal cell: 1) strengths of centrosomes’ attraction to each other and to the cell cortex have to be proportional to each other; 2) strengths of centrosomes’ attraction to kinetochores and repulsion from the chromosome arms have to be proportional to each other. We also find that three other spindle configurations emerge if these conditions are not met: a) collapsed, b) monopolar, c) multipolar spindles, and the computational screen reveal mechanical conditions for these abnormal spindles.

**Significance statement:** To segregate chromosomes, cells assemble bipolar mitotic spindle. Multiple mechanical forces generated by microtubules and molecular motors in the spindle govern the spindle architecture, but it is unclear what force balances support the bipolarity of the spindle. This problem is especially difficult and important in cancer cells, which often have multiple centrosomes that somehow are able to cluster into two spindle poles. By using stochastic energy minimization in an effective energy landscape of the spindle and computationally screening forces, we find mechanical conditions for mono-, multi- and bi-polar spindles. We predict how microtubule and motor parameters have to be regulated in mitosis in multi-centrosomal cells.

## INTRODUCTION

Many cell biological problems converge on understanding dynamic architecture of molecular machines (1), for example, mitotic spindles. During cell division, cells assemble the spindle to segregate chromosomes (2). It is crucial that the spindle is bipolar (Fig. 1), so that sister chromatids segregate to two opposite poles and end in two daughter cells. In animal cells, centrosomes (CSs) are organelles nucleating and anchoring minus ends of microtubules (MTs); a dynamic MT network spans the space between CSs and chromosomes and connect to chromosomes by plus ends. Thus, despite the fact that the exact role of CSs in the process of spindle assembly is not simple and varies between different cell types (3), more often than not, CSs organize the spindle poles (4). It is therefore not surprising that having exactly two centrosomes per mitotic cell is fundamentally related to the bipolarity of the spindle.

**FIGURE 1.**
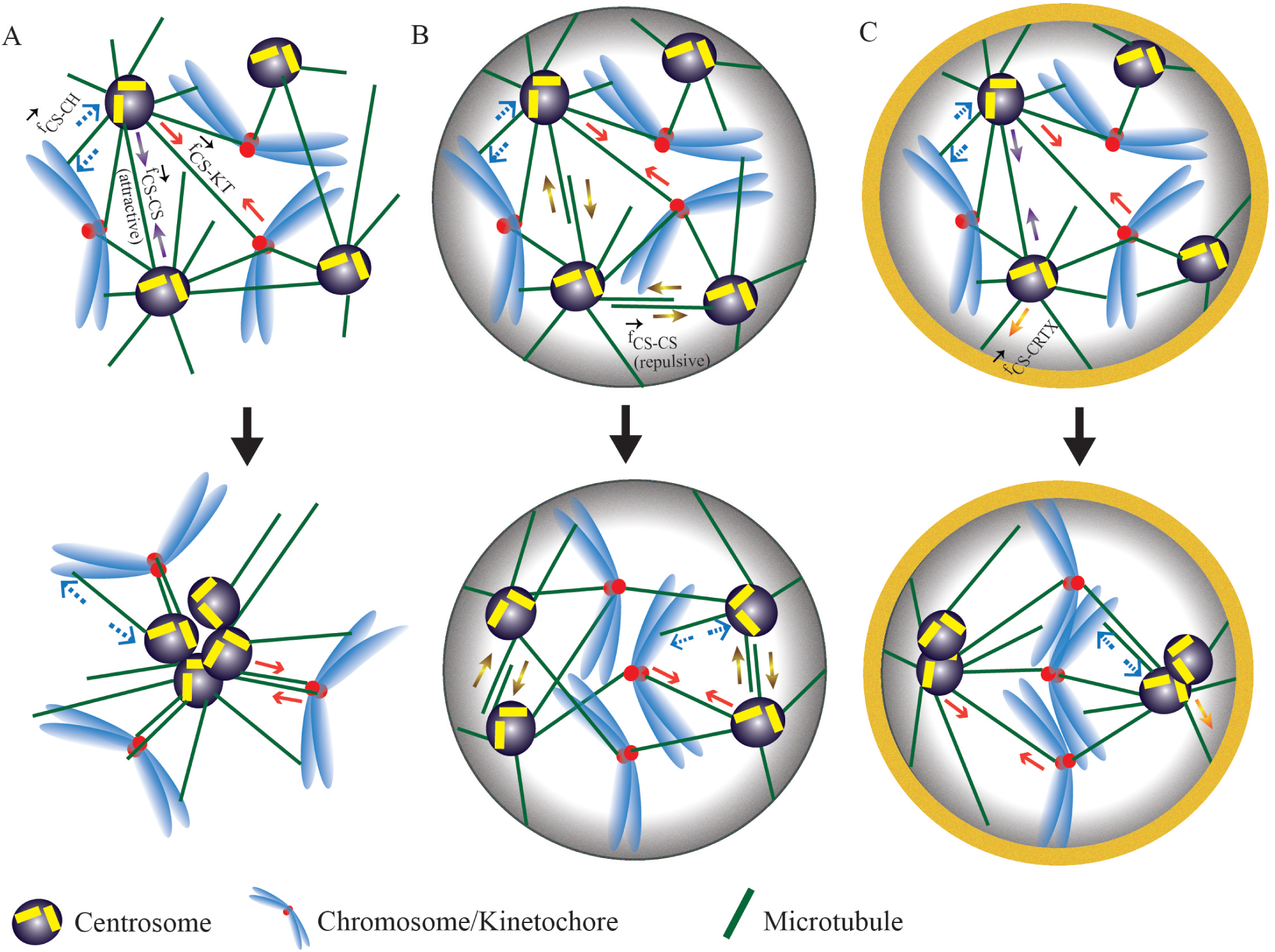
Model schematics depicting the individual forces governing the mechanics of multi-centrosomal spindles. (A) CSs aggregate into a single cluster and monopolar spindle emerges when CSs attract each other (*f*_*CS*−*CS*_), KTs (*f*_*CS*−*KT*_) and repel chromosome arms (*f*_*CS*−*CH*_). (B) In confinement of the cell volume, but without active interaction with the cortex, multipolar spindle emerges when CSs repel each other. (C) CSs aggregate into two clusters at the opposite cell poles, which chromosomes gather at the equator, creating bipolar spindle, when CSs are mutually attractive and also are attracted to the cell cortex (*f*_*CS*−*CRTX*_). The orange annular ring denotes the presence of cortical pull.

**FIGURE 2.**
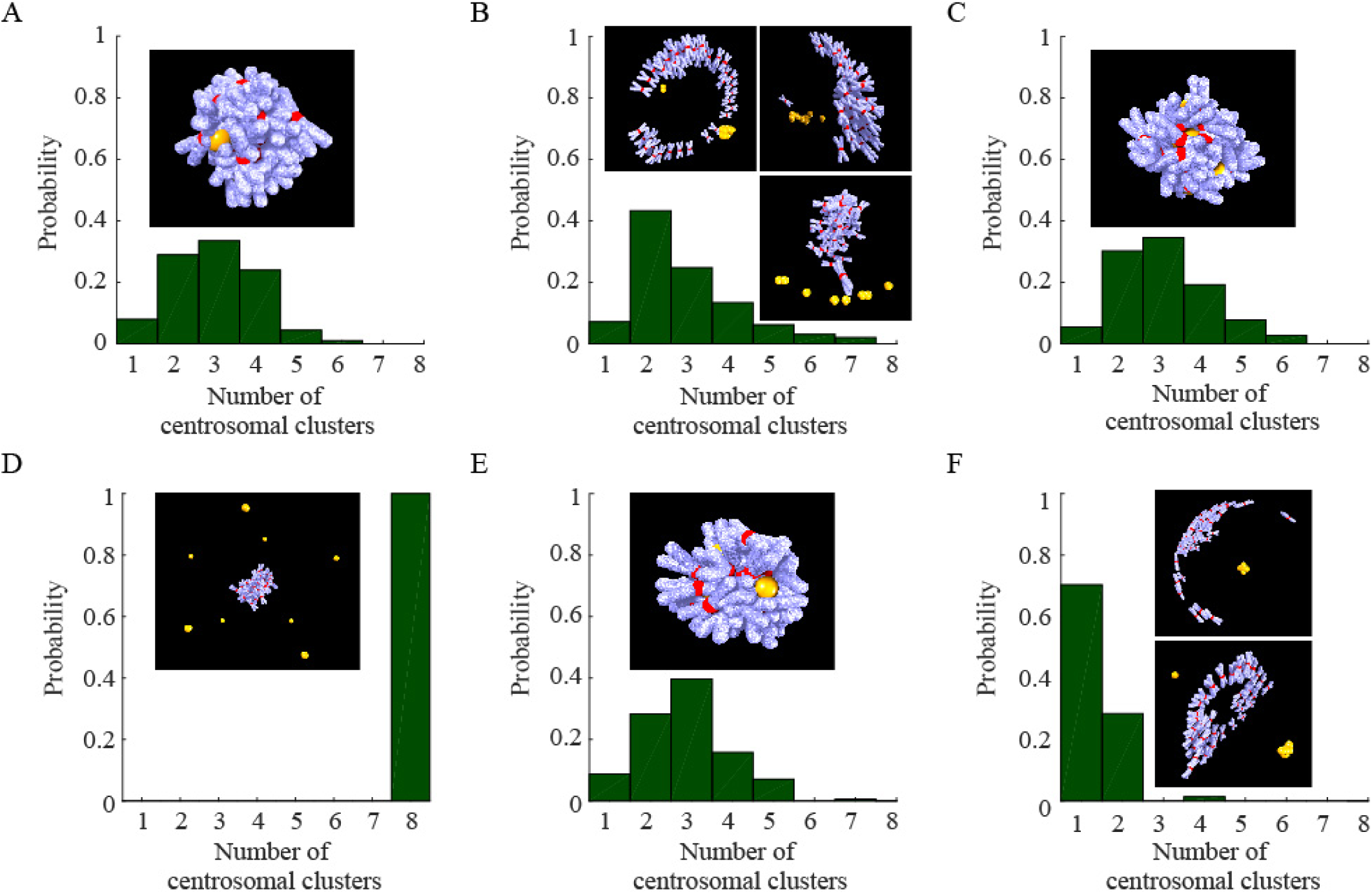
Spindle assembly in an unconfined geometry. CSs are yellow; chromosome arms are blue-and-white; KTs are red. (A) Spindle collapses - both CSs and chromosomes aggregate - in the sole presence of CS-KT attraction. (B) Both bipolar and multipolar, with a small number of monopolar spindles develop when the CS-KT attraction is combined with the CS-chromosome arm repulsion. (C) The CS-CS repulsion does not rescue the spindle from the collapse in the presence of the CS-KT attraction and absence of the CS-chromosome arm repulsion. (D) Multipolar spindles emerge in presence of CS-CS repulsion when the CS-KT attraction is combined with the CS-chromosome arm repulsion. (E) Spindle collapses when CS-KT attraction is supplemented with the CS-CS attraction. (F) Monopolar and bipolar spindles emerge in the presence of the CS-CS attraction, CS-KT attraction and CS-chromosome arm repulsion.

Normally, CSs duplicate once per cell cycle, but various perturbations can result in the accumulation of more than two CSs per cell (5, 6). Multi-CS cells are a common feature of tumors, but certain healthy cells in our body also contain extra CSs, either transiently or permanently (7). The presence of more than two CSs at the onset of mitosis has long been associated with multipolar spindle formation (4) (Fig. 1). Multiple CSs and multipolar spindles are often correlated with chromosome instability (6, 8), aneuploidy (9), erroneous merotelic attachments of chromosomes (10, 11) and other defects of chromosome segregation (12).

In recent years, several studies have shown that a process of ‘CS clustering’ -gathering multiple CSs into two groups at the opposing spindle poles - is one of the main pathways leading to the formation of the bipolar spindles in multi-CS cells (4, 5). However, not all cells appear to have equal capacity to cluster multiple CSs (11, 13); thus, understanding the CS clustering mechanisms is of great importance.

Distinct but not mutually exclusive mechanisms for spindle self-organization have been proposed. These mechanisms range from intrinsic chemical and physical, to extrinsic models (14), but the majority of the models are mechanical, rely on a balance of forces as greatest contributor to the spindle architecture (4, 14, 15). Chemical and genetic inhibition, molecular screening and micromanipulation experiments established that a great variety of molecular and geometric factors (reviewed in (4)), including minus-end molecular motors Dynein (13) and Kinesin-14 (5, 12), as well as cell shapes and forces from neighboring cells (5, 16, 17), act in the CS clustering mechanism.

The role of Dynein and Kinesin-14 led to two intuitive, but again not mutually exclusive, hypotheses: (a) minus-end molecular motors pull on MTs spanning the distance between two CSs generating effective inter-CS attraction (5) (Fig. 1). This attraction force clusters multiple CSs. (b) Dynein concentrated on a patch of the cell cortex and/or cortex contractions can reel in MTs from a few CSs bringing them together (5) (Fig. 1). However, a natural question is: why the CSs cluster into exactly two groups? Why do not such mechanisms lead to all CSs clustering into a single group causing formation of a monopolar spindle (Fig. 1)? Are these mechanisms, in fact, sufficient to prevent formation of the multipolar spindles? Are both inter-CS and CS-cortex interactions necessary? All these spindle types – monopolar (18–20), bipolar and multipolar – have been observed (5, 11, 12, 21), so what are the necessary and sufficient mechanics for the emergence of the bipolar spindle?

Measuring and quantitatively manipulating forces in the spindle is a great challenge (22, 23), and so modeling has long been a valuable tool complementing experimental research of all stages and aspects of spindle dynamics (17, 24–32). Specifically, force-balance models have been applied to reproduce the observed spindle structures, both in 1D (19, 33, 34), and in 2D geometry (19, 35). In this study, we introduce a force-balance model in realistic 3D cell geometry and computationally screen CS-CS, CS-cortex and CS-chromosome forces to answer the questions posed above.

The most straightforward approach would be the agent-based modeling (25, 27, 31, 32), in which all MTs, motors and organelles are simulated as agents obeying laws of mechanics. Despite significant advantages of such modeling, it has two drawbacks. First, the possible number of molecular motors’ combinations in various parts of the spindle is too great (reviewed in (15)), and exact mechanics of the collective motor force generation is still far from clear. Second, simulating an agent-based model in 3D is so time-consuming that screening many forces and parameters is out of question (36). Therefore, we resort to the ‘interacting particles’ model, in which each CS and chromosome is a particle interacting with other particles by pair-wise isotropic forces that depend on the distance between the interacting particles (37). Each such force results from averaged action of stochastic MTs’ and motors’ dynamics. Furthermore, rather than solving equations of motion of all particles, in this study we use energy minimization approach to determine spindle architectures corresponding to mechanical equilibria (38) in the presence of the combined forces. This approach allows us to rapidly screen the forces and delineate conditions for emergence of bipolar spindles in multi-CS cells.

We find that these conditions are: 1) strengths of centrosomes’ attraction to each other and to the cell cortex have to be proportional to each other; 2) strengths of centrosomes’ attraction to kinetochores (KTs) and repulsion from the chromosome arms have to be proportional to each other; 3) CS attraction to the cortex has to be short-ranged, while CS interactions with the chromosomes have to be on the scale of the cell size. We also find that three other spindle configurations emerge if these conditions are not met: a) collapsed, b) monopolar, c) multipolar spindles (Fig. 1), and the computational screen reveal mechanical conditions for these abnormal spindles. The model correctly reproduces the ‘doughnut’-like distribution of the chromosomes in the spindle and highlights the importance of the initial conditions for the spindle development.

## METHODS

There are two types of interacting bodies in the model: CSs and chromosomes; in addition, the CSs interact with the cell cortex. The CSs and chromosomes reside in the cell, which is a prolate ellipsoid, and the cortex ix the 2D surface/boundary of the cell. The bodies interact with four types of forces depicted in Fig. 1. Each force is a so-called ‘conservative’ force, and so a potential energy corresponds to each pairwise interaction. The sum of all these pairwise energies constitute the total potential energy of the system.

### Interactions between pairs of CSs

Each CS is a center of a MT aster with MTs undergoing the dynamic instability (39, 40). When a MT growing from one CS reaches another CS, or overlaps with a MTs growing from that other CS, an effectively attractive or repulsive (depending on the motor types) force, *f*_*CS*−*CS*_, can be generated by molecular motors interacting with these CSs and MTs (Fig. 1). Using conventional assumptions of a large number of isotropically distributed MTs (19,41–43), we introduce the expressions for *f*_*CS*−*CS*_, and for respective potential energy of moving one CS from distance *r*_1_ to *r*_2_ away from another CS:

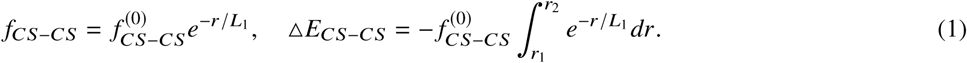

Here 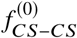 is the force between a pair of proximal CSs, *r* is the distance between CSs and *L*_1_ is the spatial range of the force. Here we utilize one of the most simple and frequently used exponentially decreasing spatial dependence of the force; in the Supporting Information we explore other spatial dependencies.

### Interaction between a CS and a KT

There are two interactions between the CSs and chromosomes: CS-KT interactions and interactions of CSs with chromosome arms. The so-called K-fibers – MT bundles – connect the CS-KT pairs, and molecular motors on the KTs and K-fibers generate net CS-KT attraction force (44) (Fig. 1). We use the simplest assumption that this force is constant, length-independent (19, 45): 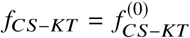. In certain cases, for example when we simulate the CSs and chromosomes not restricted by the cell boundary, we use a cutoff distance for the CS-KT force, such that the force is constant below the cutoff and equal to zero above the cutoff. Corresponding potential energy is:

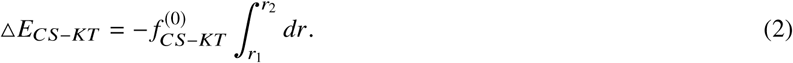

### Interaction between a CS and a chromosome arm

Another interaction is a repulsion between a CS and chromosome arms *f*_*CS*−*CH*_ (Fig. 1). This force originates both from MT polymerization forces and from kinesins on the arms interacting with the MT plus ends, and it decreases with distance (46–50). We use the following expression for this force and respective potential energy:

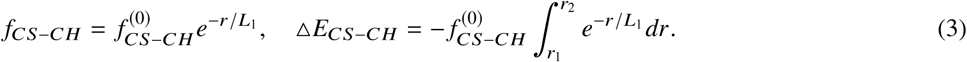

Note that all three pairwise forces introduced above are between point-like bodies and they are directed along the vectors connecting respective pairs of bodies. CS-KT and CS-chromosome arm forces between a given CS and a given chromosome are directed the same, as KT and chromosome arms of a given chromosome is the same point-like body in the model. Note also that the CS-CS and CS-chromosome arm forces are assumed to have the same ranges; the underlying assumption is that average lengths of MTs connecting the CSs and extending from the CSs to the chromosome arms are the same.

### Interaction between a CS and the cortex

MTs from the CSs reach the cortex on the inner cell boundary, and Dynein motors on the cortex pull on these MTs generating the attraction (51–53). We use the following expression for this force:

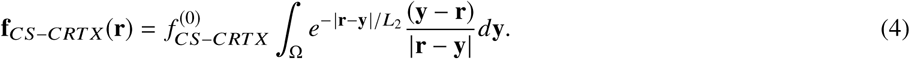

Here, *L*_2_ is the average range of the attraction to the cortex; **y** is the coordinate of a point on the cell surface; **r** is the 3D coordinate of the CS. The integration is over the cell surface **Ω**; in the simulations, the surface is approximated by a discrete grid, as explained below, and the integral becomes the sum over the grid’s nodes. The corresponding energy of moving the CS from point **r**_1_ to point **r**_2_ is:

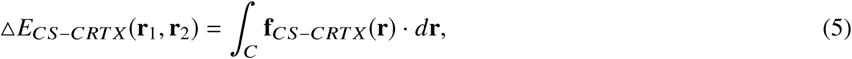

where *C* is a segment starting at **r**_1_ and ending at **r**_2_.

### Numerical implementation and energy minimization

In principle, we could have solved equations of motion for the CSs and chromosomes as follows: *z*_*i*_*d***r**_*i*_/*dt* = **G**(**r**_*i*_) + ∑ **F**(**r**_*i*_ − **r**_*j*_), where *z*_*i*_ is the drag coefficient, **r**_*i*_ is the position of the *i*^*th*^ body, **r**_*j*_ are positions of all other bodies, **F**(·) are the pair-wise forces, and **G**(·) is the force from the cortex. However, the rules of movement are in fact unknown, and in any case we are only interested in the mechanical equilibria. Thus, we introduce the total system’s mechanical energy: *E* = *P*_1_ (**r**_*i*_) + ∑ *P*_2_ (**r**_*i*_ − **r**_*j*_), where *P*_1_ and *P*_2_ are the potential energies corresponding to the cortex and other forces introduced above, and search for the geometric configurations minimizing this energy. Such approach was considered and justified in applications to macroscopic biological problems (54, 55). Considering cell states as minima in the energy landscapes is gaining popularity (56); note that we consider a well defined mechanical energy, as in (55), rather than less concrete total energy which is not easy to define in the open system of a cell. Note also that computer simulations are necessary to explore the model energy landscapes, because mechanical equilibria of models even simpler than ours can be very complex (57).

In most of the the simulations, we consider 8 CSs and 46 chromosomes. For simplicity, both CSs and chromosomes are represented by point-like objects. There is a steric repulsion between pairs of chromosomes that scales as inverse square mutual distance and is on when the mutual distance falls below 2 units of the numerical grid. The steric repulsion between two CSs or between CSs and chromosomes is not considered explicitly, but one body is prohibited from moving onto a site, which is already occupied by any other body. When the spindle is simulated within the cell, CSs and chromosomes are not allowed to pass through the cell boundary.

In simulations without the cell, we chose a large cubic lattice of size 120×120×120 *μm*^3^ and all the CSs and chromosomes were distributed randomly within a small sphere at the center of this lattice at the onset of the simulation. In the cellular geometry, a prolate ellipsoid with semi-axes 20 *μm*, 15 *μm* and 15 *μm* is introduced, and the chromosomes and CSs are initially randomly placed within the confinement. The cell volume is fragmented into a 3D cubic lattice grid. The grid size is 1 *μ*m for all cases. The ellipsoid of the cortex is discretized into a finite number of approximately equidistant nodes with the grid size similar to that in the cell volume.

The configuration of the system is updated using Monte Carlo method in the following manner:

1. A body (CS or chromosome) is selected randomly.
2. If it is a CS, move it to a vacant neighboring site chosen randomly and calculate the respective energy change Δ*E* for moving it from initial position *r*_1_ (node from which the CS moved) to a neighboring position *r*_2_ (node to which the CS moved) by adding Δ*E*_*CS*−*CS*_, Δ*E*_*CS*−*CH*_, Δ*E*_*CS*−*KT*_ and Δ*E*_*CS*−*CRTX*_ (the latter – if the spindle is in the cell).
3. If it is a chromosome, do the same, but the energy change is calculated by adding Δ*E*_*CS*−*CH*_ and Δ*E*_*CS*−*KT*_ only.
4. The move is accepted if Δ*E* ≤ 0, else the move is accepted with the probability *p* = *e*^*−βΔE*^ (Boltzmann weight), where *β* is the inverse of an effective thermal energy (temperature) required to update the configuration (58). Effective temperature, 1/*β*, was chosen from numerical trials. As we report in the Supporting Material, the energy differences and the energy barriers between local minima in the spindle energy landscape are on the order of 100 pN×*μ*m. When the chosen temperature is much lower than this characteristic value, the spindle in the simulations is frozen in a minimum near an initial configuration and does not evolve. When the chosen temperature is much higher, the spindle never settles in any energy minimum and keeps jumping randomly between configurations. Thus, we used *β* = 100 pN×*μ*m, which is on the order of the the energy barriers between local minima in the spindle energy landscape.
5. The simulations are carried out until a stable equilibrium configuration is achieved. More precisely, we calculate the energies of the system during 1000 time-steps, by the end of which the system is normally not evolving, and energy is not changing. Then we repeat the same procedure for the next 1000 time-steps as well. If the energy difference between the earlier average at the end of the first run, and the latter average is less than a threshold of (∼ ± 1 pN ×*μ*m), we conclude that the system has attained equilibrium. The equilibrium statistics have been obtained by averaging over ∼ 100-200 random initial configurations. In the equilibrium configuration, if 2 or more CSs are within a distance *L*_*merge*_ (which is equal to 1.5 *μ*m, corresponding to when two CSs are in the neighboring nodes of the numerical grid), they are considered to be clustered.

The model parameters are listed in Table S1 in the Supporting Material. The code for the Monte Carlo simulation written in C can be downloaded from the in the Supporting Material. The data analysis and plotting have been carried out in MATLAB. A single simulation run takes about 40 min of real time until a mechanical equilibrium is achieved (in Intel(R) Xeon(R) CPU having clock speed 2.20 GHz, RAM 50 GB).

## RESULTS

We simulated the model with various combinations of the four principal forces; detailed results are reported below and depicted in Fig. 2-4. In brief, first, we simulated the spindle in the absence of the cell boundary, essentially considering the spindle in an unconfined space. Bipolar spindles do emerge in these simulations, but not robustly (Fig. 2). Then, we introduced the cell boundary and cortex and simulated either non-interacting, or mutually repulsive, CSs, and again did not detect the robust bipolarity (Fig. 3). We found that the bipolar spindle emerges robustly when the CSs are attracted to each other and to the cortex and interact with both KTs and chromosome arms (Fig. 4). In the end, we explore dependence of the spindle architecture on mechanical and structural parameters and initial conditions (Fig. 5). Note that in the figures and movies, the chromosome arms shown in the computational figures are there for the visual effect only, as in the models the chromosomes are the point-like bodies. The statistics of the simulated spindles is summarized in Table S2 in the Supporting Material.

**FIGURE 3.**
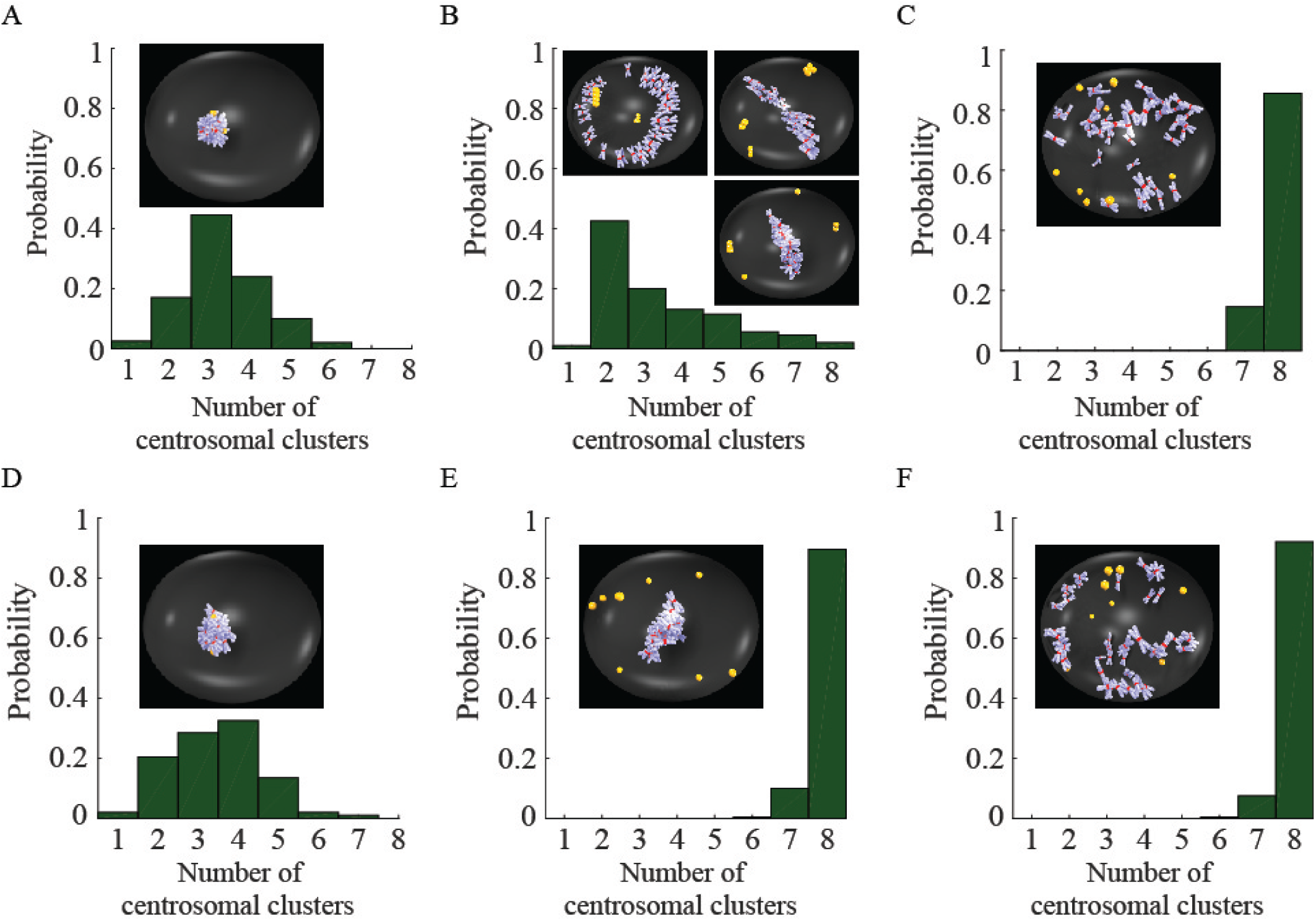
Spindle assembly in the presence of the CS-cortex attraction without CS-CS attraction. CSs are yellow; chromosome arms are blue-and-white; KTs are red. (A) Collapsed spindle in the presence of the CS-KT attraction and in the absence of the inter-CS interaction. (B) Non-robust emergence of the bipolar spindle in the presence of the CS-KT attraction, CS-chromosome arm repulsion and in the absence of the inter-CS interaction. (C) Multipolar spindles develop under the sole influence of CS-chromosome arm repulsion. (D) Collapsed spindle when the CS-KT attraction is supplemented by the inter-CS repulsion. (E) Multipolar spindles develop when the CS-KT attraction and CS-chromosome arm repulsion are supplemented by the inter-CS repulsion. (F) Multipolar spindles develop when the CS-chromosome arm repulsion is combined with inter-CS repulsion.

**FIGURE 4.**
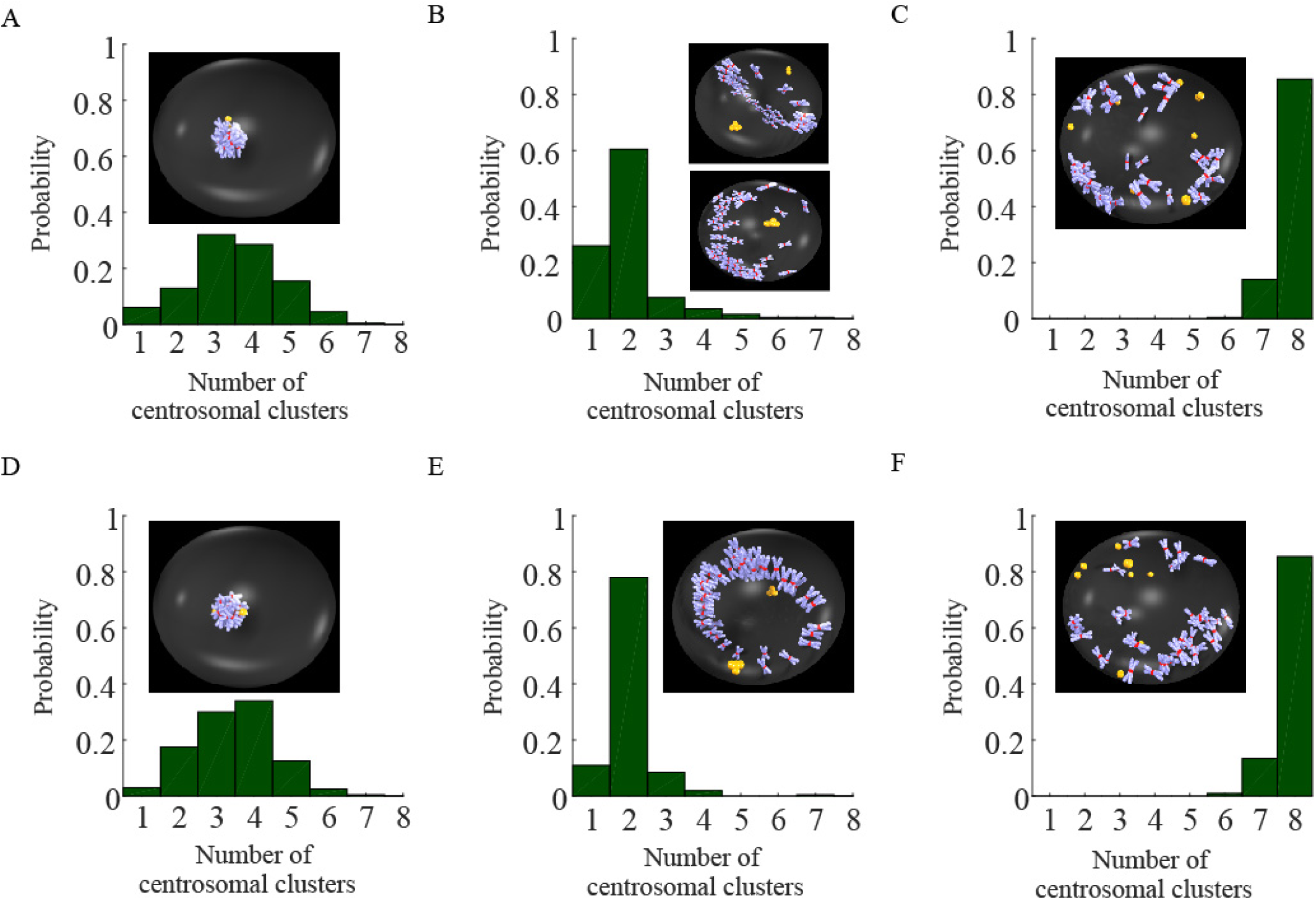
Spindle assembly in the presence of the CS attraction to each other and to the cortex. In all simulations reported here, the CS-CS attraction is present. CSs are yellow; chromosome arms are blue-and-white; KTs are red. (A) Spindles collapse when CSs are attracted to KTs without the CS-cortex attraction. (B) Bipolar spindles emerge more robustly with combination of the CS-KT attraction and CS-chromosome arm repulsion in the absence of the CS-cortex attraction. (C) Multipolar spindles develop when only the CS-chromosome arm repulsion, but not CS-KT attraction, act in the absence of the CS-cortex attraction. (D) Spindles collapse when CSs are attracted to KTs and to the cortex. (E) Bipolar spindles evolve most robustly when the CS-cortex attraction, CS-KT attraction, CS-chromosome arm repulsion and CS-CS attraction are combined. (F) Multipolar spindles develop when the CS-cortex attraction, CS-chromosome arm repulsion and CS-CS attraction are combined, but the CS-KT attraction is absent.

**FIGURE 5.**
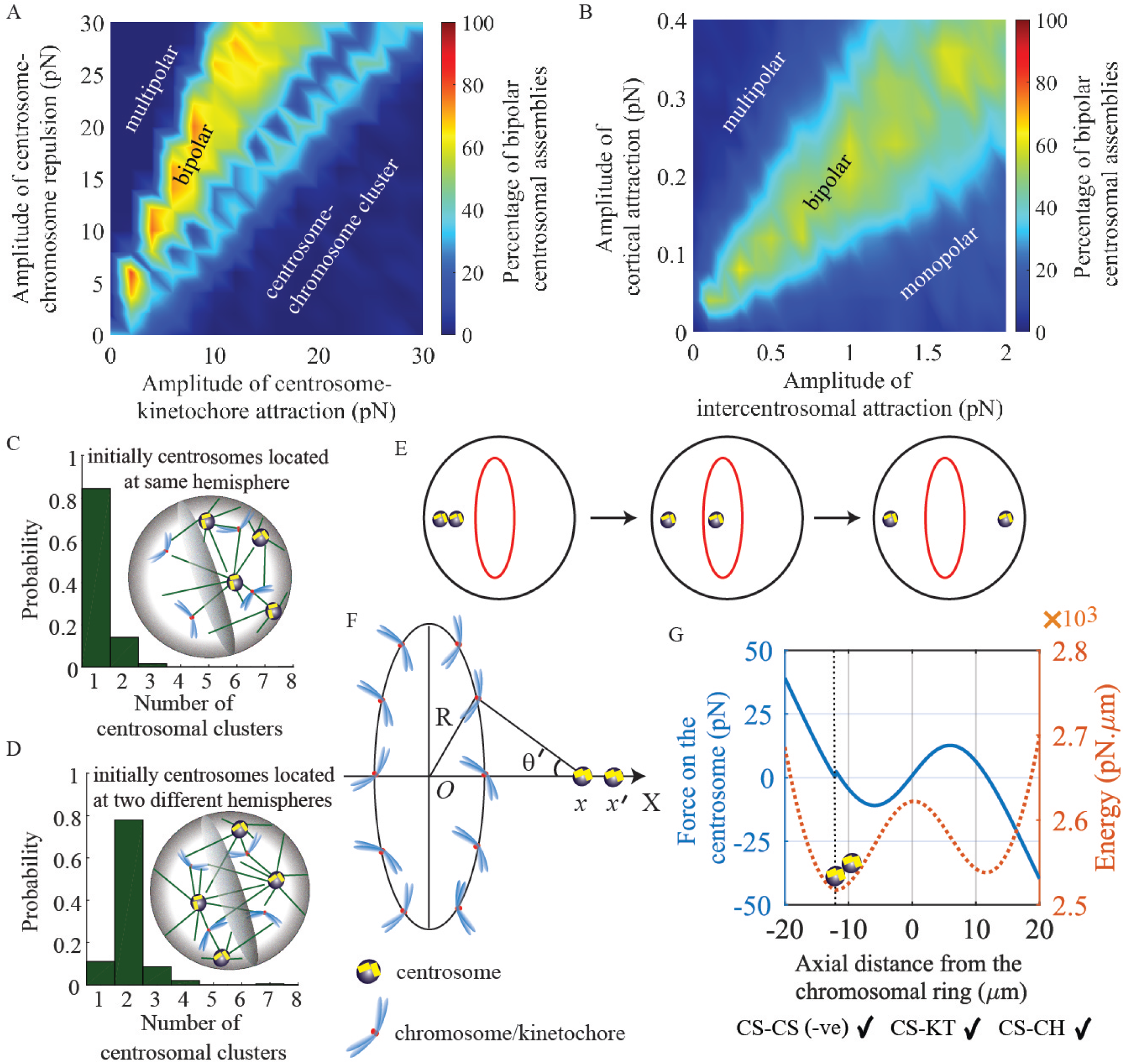
Bipolar CS clustering is sensitive to mechanical and structural parameters and initial conditions. In all simulations, forces or parameters other than varied ones are same as those used in simulations for Fig. 4E. (A) Percentage of the emerged bipolar spindle as the function of the amplitudes of the CS-KT attraction and CS-chromosome arm repulsion. (B) Percentage of the emerged bipolar spindle as the function of the amplitudes of the CS-cortex attraction and CS-CS attraction. (C-D) Relative frequencies of the CS cluster numbers when eight CSs are initially localized in the same (C) and different (D) hemispheres. (E) Scheme of the numerical experiment: the chromosomes are uniformly distributed along the circumference at the cell equator, one CS is placed to the cell pole, while another CS is gradually moved from that pole to the other one. (F) Coordinates and angles of the numerical experiment. (G) Force acting on the moving CS in the numerical experiment and respective energy of the system as function of the inter-CS distance.

### Bipolar spindles do not assemble robustly in unconfined geometry

We first explored the emerging spindle architecture without the complexity of interacting with the cortex or being restricted by the cell geometry. If only one type of force, the CS-KT attraction, is present, the spindle collapses (Fig. 2A, inset): all CSs and chromosomes aggregate together. This result is easy to understand: attraction brings the chromosomes to the CSs, while this very same attraction brings other CSs to the aggregate. Note that the emerging number of the CS clusters varies (Fig. 2), depending on the initial conditions. All these clusters are very close to each other; they do not merge due to the steric repulsion (volume exclusion) effect; effective energy barrier of pushing crowded chromosomes between CSs out of the way is too high.

Adding the CS-chromosome arm repulsion to the CS-KT attraction rescues the spindle: ∼40% of the emerged spindles are bipolar (Fig. 2B). The repulsion prevents the CSs and the chromosomes from collapsing onto each other. Indeed, because the CS-KT force is attractive and distance-independent, while the CS-chromosomal arm force is repulsive, decreases with distance and is greater than CS-KT attraction at small distances, there is a stable mechanical equilibrium such that the CSs and chromosomes tend to rest at a finite distance from each other. Note that in the emerged bipolar spindle at the inset of Fig. 2B, the chromosomes gather into a ‘doughnut’, and CSs are in two clusters, equidistant from the chromosomes. However, the bipolar geometry is not robust: many multipolar or monopolar spindles emerge, with chromosomes aggregating into one complex-shaped manifold, and CSs again equidistant from this manifold, but not bi-clustered (Fig. 2B inset).

If, instead of the CS-chromosome arm repulsion, only CS-CS repulsion is combined with the CS-KT attraction, the collapsed spindles are not rescued (Fig. 2C). The reason is that the total energies of both CS-KT attraction and CS-chromosome arm repulsion scale as the product of the numbers of the CSs and chromosomes, while the energy of the CS-CS interactions scale with the square of the CS number. The number of CSs is much smaller (more than five-fold) than the number of the chromosomes, and the energy of attraction in this case overwhelms the repulsive energy.

If we add the CS-CS repulsion to the interactions of the CSs, KTs and chromosome arms in the case depicted in Fig. 2B, the CSs tend to stay at a constant equilibrium distance from the chromosomes, and also as far as possible from each other. This leads to the multipolar spindles, in which all chromosomes aggregate at the center, while the CSs are scattered uniformly across a spherical surface centered at the chromosomal crowd (Fig. 2D).

Finally, we turned to the case when CSs are mutually attractive. As expected, when only attractive interactions - CSs attract each other and KTs - are present, the spindle collapses (Fig. 2E).

When we add the CS-CS attraction to the case of the Fig. 2B - combined CS-KT attraction and CS-chromosome arm repulsion - the multipolar spindles disappear, because now the CSs attract each other, and majority of the spindles are monopolar, with a single CS cluster at the center and chromosomes scattered at a spherical surface centered at the CS cluster (Fig. 2F, Movie M1). However, some bipolar spindles also emerge, with the architecture similar to those in the absence of the CS-CS attraction. The reason is that the chromosomal ‘doughnut’ in the middle repels the CS clusters at both sides away from the middle creating the energy barriers, which the CS-CS attraction cannot overcome.

Lastly, in the absence of the CS-KT attraction, repulsion of the CSs from the chromosome arms keeps the spindles multipolar even in the presence of the CS-CS attraction (Table S2). The bottom line is that the combination of the CS-KT attraction with CS-chromosome arm repulsion helps the bipolar spindles to emerge, but the bipolar architecture is not robust without interactions with the cell boundary.

### Without inter-CS attraction, bipolarity does not emerge in cell confinement

As the bipolar spindle architecture cannot be achieved without interactions with the cell boundary, we next turned to testing the spindle mechanical equilibria in the cell confinement. We found that, other than in the case of the CS-CS attraction, which we describe below, just confinement of the spindle in the cell has the same effect as the CS-cortex attraction (compare Fig. 3 and Fig. S1). Therefore, here we only discuss the results in the presence of the CS-cortex attraction.

We find that spindles collapse in the presence of the CS-KT attraction and in the absence of the inter-CS interaction (Fig. 3A), same as in the unconfined case. The only effect of the CS-cortex attraction is that the collapsed spindle is located near the cortex. The collapse is not rescued by addition of the CS-CS repulsion (Fig. 3D), same as in the unconfined case.

When we combine CS-KT attraction and CS-chromosome arm repulsion, the bipolar spindle geometry develops (Fig. 3B), but not robustly: only ∼ 40% of the spindles are bipolar. Thus, the presence of the attraction to the cortex does not make the bipolarity robust (compare to Fig. 2B) in the absence of the CS-CS interactions. If we make the CS-CS interactions repulsive, then, as expected, all emerging spindles become multipolar (Fig. 3E, Movie M2).

Lastly, if only CS-chromosome arm repulsion is present, then, predictably and similar to the unconfined case, the multipolar spindles emerge either in the absence (Fig. 3C) or presence (Fig. 3F) of the CS-CS repulsion. To conclude, the confinement and attraction of the CSs to the cortex does not make the bipolar spindle architecture robust.

### Combination of the CS attraction to the cortex, to each other and to KTs with repulsion from the chromosomal arms makes the bipolar CS clustering robust

We finally tested the few remaining cases in which CS-CS attraction was present. We started with three cases on the spindle confined in the cell but the CSs are not attracted to the cortex, and then explored the effect of the CS attraction to the cortex. Same as in the previously considered cases, in the absence of the CS-chromosome arm repulsion, the spindles collapsed in confinement (Fig. 4A) and in the presence of the CS-cortex attraction (Fig. 4D). Similarly, as in the previously considered cases, in the absence of the CS-KT attraction, multipolar spindles emerged in confinement (Fig. 4C) and in the presence of the CS-cortex attraction (Fig. 4F).

Eventually, we explored the combination of the CS-KT attraction, CS-chromosome arm repulsion and CS-CS attraction, which promotes the bipolarity non-robustly in the unconfined space (Fig. 2F). We found that in confinement the bipolarity becomes more robust: ∼ 60% of the evolved spindles were bipolar (Fig. 4B), compared to ∼ 40% in the unconfined case. Even better result is achieved in the presence of the active CS-cortex attraction: in ∼ 80% of cases, the CSs aggregated into two clusters (Fig. 4E, Movie M3) creating the familiar ‘doughnut’ chromosome structure. The more robust bipolar CS clustering in this last case is due to attraction of the evolving opposite CS clusters to the cortex that prevents them from falling onto each other, assisted by the repulsion from the chromosomal ‘doughnut’.

### Bipolar CS clustering is sensitive to mechanical and structural parameters and initial conditions

#### Sensitivity to model parameters

To test how sensitive is the spindle architecture to the model parameters, we varied single and pairs of parameters, keeping the rest of the parameters equal to the base values listed in Table S1, and repeated the simulations. Note that the base parameter values predict the robust bipolar spindles reported in Fig. 4E. We found that two force balances have to hold for the bipolarity. First, CS-KT attraction strength has to be proportional to the CS repulsion from the chromosome arms (Fig. 5A); if the repulsion is weaker/stronger than the attraction, then the spindle collapses / becomes multipolar. Second, the strength of the CS-CS attraction has to be proportional to the CS-cortex attraction (Fig. 5B); if the interaction with the cortex is weaker/stronger than the CS-CS attraction, then the spindle collapses / becomes multipolar. Sensitivity of the spindle architecture to the individual forces’ strengths is further demonstrated in Fig. S2B,C.

Next, we investigated sensitivity to varying force ranges. Recall that the CS-KT attraction is distance-independent in the model, and so there are three force ranges in the model – for the CS-chromosome arm repulsion and for the CS-CS and CS-cortex attractions. We varied these ranges and found that the range for the CS-chromosome arm repulsion has to be on the order of the cell radius (Fig. S2A) for 2 CS clusters to emerge. The reason is rather simple: if this force range is too small, the repulsion is effectively too weak, and the spindle collapses. In the other limit, the repulsion is too strong pushing the CSs into the cortex too hard for the CS-CS attraction to cluster them. On the other hand, the bipolarity is not very sensitive to the range of the CS-CS attraction (Fig. S2A). Finally, we also varied the range of the CS-cortex attractions and found that this length has to be much smaller than the cell size in order for the bipolar spindles to emerge (Fig. S4C). This can be explained if one considers the integrated effect of the whole cortex on one CS. It was demonstrated in a number of models (59) that if the range of the attraction to the cortex is long enough, then effectively the force on the CS becomes centering. In that case all CSs are pulled to the cell center creating the monopolar spindle.

Lastly, we repeated all simulations for the normal cells with two CSs. As can be seen from Fig. S3C,D, proper bipolar architectures with the chromosomes between two segregated CSs emerge most robustly when there are either no CS-KT interactions, or when CSs *repel* each other. This makes easy physical sense: two CSs segregate better when they repel, not attract each other. The same conclusion was reached in a number of previous models (19, 33, 34). Other numerical tests are described in the Supporting Material.

#### Sensitivity to the initial conditions

How important is it where the CSs are placed initially, at the end of prophase – early prometaphase? We simulated two scenarios: first, the CSs were scattered uniformly across the whole cell; second, the CSs were located with uniform probability only in half (one hemisphere) of the cell. Fig. 5C,D shows that in the first case the bipolar spindle evolves, while in the second case – monopolar one. The explanation is that if initially the CSs are close together, their attraction overcomes the repulsion of a small number of the chromosome arms between the individual CSs. Thus, the proper spindle architecture is sensitive not only to the mechanical parameters, but also to the initial conditions. The same conclusion holds for the cell with two CSs (Fig. S4E,F).

#### Mechanical energy landscape of the bipolar spindle

To have a better understanding of the spindle bipolarity, we considered a simple analytical model, mimicking the proper spindle geometry: the chromosomes arranged uniformly in an annular-shaped ring of inner radius *r*_*a*_ and outer radius *r*_*b*_ (Table S1) within a spherical cell of radius *R*_*cell*_ (Fig. 5E-5F). Two CSs are positioned on the axis through the center of the ring, perpendicular to the plane of the ring. In the presence of the CS-chromosome arm repulsion and CS-KT and CS-CS attraction, we calculated (Supporting Material) the configuration’s energy as function of the CS-CS distance, when one CS is at the left from the chromosomal ring, and another moves to the right along the axis. Fig. S4H shows that when there is no CS-CS interaction, there is a double-well energy profile for this spindle, with equally deep energy wells. Each well corresponds to a CS being at a mechanically equilibrium distance from the chromosomal ring, at which the CS-KT and CS-chromosome arm forces are balanced to zero. As the CSs do not interact in this case, the mono- and bi-polar spindle have the same energies, with a significant energy barrier between them, due to the difficulty of pushing the CS through the chromosomal ring. If we add the CS-CS attraction, the double-well energy profile changes, with the well corresponding to the monopolar spindle becoming deeper than the well corresponding to the bipolar spindle (Fig 5G). This explains why the monopolar spindles are the most stable (see the Supporting Material); however, the bipolar spindle is very stable as well, because the energy barrier between them remains significant. Lastly, this result also illustrate the predicted sensitivity to the initial conditions: If we start with both CSs to the left from the chromosomes, they both ‘fall’ into the same energy well making the monopolar spindle, but if initially the CSs are at the opposite sides of the chromosomes, they each ‘fall’ into its own energy well making the bipolar spindle.

## DISCUSSION

In this study we examined computationally mechanical requirements for the bipolar CS clustering and reached the following conclusions. Four types of forces are necessary for the bipolar spindle architecture: CS attraction to each other, to the KTs and to the cortex, and CS repulsion from the chromosome arms. In order for these forces to be sufficient for the bipolarity, four further conditions have to be met: CS attraction to each other needs to be proportional to the attraction to the cell cortex; CS attraction to the KTs has to be proportional to the repulsion from the chromosome arms; interactions of the CSs with both KTs and chromosome arms have to be on the scale of the cell size; CS attraction to the cortex has to be short-ranged. Physical explanation of the bipolarity under these conditions is this: the balance of the constant attraction to the KTs and repulsion from the chromosome arms decreasing with distance on the scale of the cell places CSs at certain equilibrium distance from the chromosomes, and this equilibrium distance is on the order of the cell size. If CSs attract each other *and* attraction to the cortex is strong *near the cortex*, then, providing that initially the CSs are scattered around the cell, proximal CSs start clustering into greater and greater clusters, and at some point there is one cluster in one hemisphere of the cell, and another one – in another hemisphere. Most of the chromosomes trying to stay away from both CS clusters are in the middle, creating a barrier for the CS clusters; in addition, the cortex pulls two CS clusters into the opposite directions, stabilizing the bipolarity. If the attraction to cortex is global, nothing can prevent the collapse of the two clusters.

The simulations, interestingly, successfully reproduce the characteristic ‘doughnut-like’ spatial organization of chromosomes in prometaphase (60). The physical explanation is that for chromosomes, staying away from two opposite CS clusters at the cell poles, the doughnut around the cell equator is the place where all chromosomes are at equilibrium distance from the cell poles.

If CSs attract each other but the interactions with the cortex are weak, CSs cluster into one group at the cell center, and chromosomes spread over the cortical surface, and the monopolar spindle emerges. If, on the other hand, the interactions with the cortex are weak, but CSs repel each other (or do not interact with each other), the chromosomes gather at the center, while CSs spread randomly over the cortex, making the multipolar spindle. Also, if there is no connection between the KTs and CSs, the spindle is multipolar. Both mono- and multipolar spindles were observed. Finally, without the repulsion from the chromosome arms, the whole spindle collapses; this, to our knowledge, was never observed, perhaps because there is always some repulsion from the chromosome arms.

Our model predictions are in general qualitative agreement with the published data: minus-end motors Dynein (13) and Kinesin-14 (5, 12), normally associated with attractive forces in multiple models, are needed for CS clustering, suggesting that both attractions of CSs to the cortex and each other are necessary. Higher activity of plus-end Kinesin-5 motor, normally associated with repulsive forces in multiple models, diminishes the clustering effect (61). Perturbations of chromokinesins that contribute to the repulsion of the CSs from the chromosome arms leads to spindle defects (62, 63). A number of studies suggested that MTs, MT associated proteins and motors, affecting not only cortex and CS-CS forces but also CS-KT forces, contribute to clustering (5, 13, 64, 65). This is in agreement with the prediction that magnitude of the CS-KT attraction affects the clustering effect (Fig. 5S, Fig. S2B). A few studies demonstrated that while inactivation of one type of motor disrupts spindle morphology, the phenotype can be rescued by simultaneous inactivation of another, opposing, motor (66–68). Such pairwise inactivation of opposite motors is likely to diminish two opposing forces, while keeping their ratio unchanged, and so stability of the spindle morphology under such double perturbations is consistent with the model. CSs that lack chromosomes between them do not form a stable spindle-like MT array (69), in agreement with the model. The predicted sensitivity to the initial conditions - defects in the spindle architecture when the CSs are initially too close to each other - was documented in (70).

It was shown that if interaction of CSs with the cortex is not uniform, but rather CSs attract to local patches on the cortex, which is the case for cells with anisotropic and heterogeneous adhesion patterns, then multipolarity of the spindle is enhanced (5, 16, 17). Though we have not simulated such situation explicitly (this is a worthy future effort), qualitatively it is clear that more than two localized adhesive patches would attract locally smaller CS clusters preventing them from merging into bigger ones. The model also can be expanded to explore non-trivial role of cell shape in positioning and shaping the spindle (71).

It is thought-provoking to explore progression of the spindle through stages of mitosis from the model’s point of view. There are no MT-KT connections in early prometaphase, so the model says that at this stage the spindles in multi-CS cells are multipolar. Later, when enough MT-KT connections are made, CS clustering follows. Interestingly, multipolar spindles in early prometaphase and CS clusters in late prometaphase actually were observed in Drosophila SakOE neuroblast spindles (72), in agreement with the prediction. This sequence of events, in fact, could enforce initial conditions benefitting later bipolarity by pushing apart CSs at the early stage preventing all of them ending up in the same hemisphere of the cell.

Many various combinations of molecular motors can generate forces necessary for the bipolar CS clustering. To explore relevant molecular pathways deeper, agent based modeling will be needed. Such stochastic modeling will also circumvent the problem of choosing ‘temperature’ in our energy minimization approach: the temperature associated with the heat bath in the Monte Carlo simulation algorithm are manifestations of stochastic fluctuations arising from the MT dynamic instability and finite numbers and stochastic dynamics of the motors. Besides stochastic effects, other aspects of the spindle mechanics will have to be considered to make the model less simplistic. Those include influence of merotelic errors of assembly on the force balance (25), space-dependent motor regulation (15), viscoelastic properties of the spindle (23), and role of the cortex flow in bringing CSs together (73), not to mention biophysical details of the magnitudes and spatial dependencies of the forces in the spindle that remain under-researched (22, 47, 74).

The predicted conditions for the spindle bipolarity in multi-CS cells are stringent, including proportionality of two pairs of forces, as well as strict limits on the ranges of these forces. (It is worth noting that proportionality of pairs of opposing forces was also predicted for length regulation of spindles in early *Drosophila* embryo (34).) This could mean that the predicted mechanical CS clustering mechanism is not very robust. However, there could be mechanochemical feedbacks robustly regulating the mechanical parameters to satisfy exactly these conditions. Another possibility is that additional mechanisms are in place to make the bipolar CS clustering more robust. One distinct possibility is that CSs are not equal: our model suggests that the robust bipolar spindle emerges in 2-CS cell when these CSs repel each other. If these two CSs are made ‘dominant’, while other CSs are ‘inactivated’, or if CSs are attracted not individually to each other, but rather to poles of a dominant interpolar MT bundle (both ideas are reviewed in (4)), then the bipolar spindle could emerge more robustly. Yet another factor that could enhance the robustness is a non-isotropic CS-cortex interaction. Last, but not least, there is no ‘the’ spindle: different cell types evolved to use differently the common molecular tool box to build bipolar spindles (75). Comparing design principles behind different mechanisms to cluster multiple CSs in mitosis is one of the future goals.

## CONCLUSION

To assemble the bipolar spindle in a multi-centrosomal cell, the centrosomes have to attract proportionally to each other and to the cell cortex; also centrosomes need to have proportional attraction to the kinetochores and repulsion from the chromosome arms. In addition, the centrosome-cortex interaction has to be short-ranged, while the ranges of the centrosome-kinetochore and centrosome-chromosome arm interactions have to be comparable with the cell size. Without centrosome-chromosome arm repulsion, the spindles collapse. Without centrosomes attracting to the cortex, monopolar spindles evolve if centrosomes mutually attract, and multipolar spindles – if centrosomes do not interact or repel each other. Spindle architecture is sensitive to initial conditions.

## AUTHOR CONTRIBUTIONS

A.M. and R.P. conceived and directed the study with help from A.K.; S.C., A.M. and R.P. wrote the manuscript with help from A.K.; S.C., A.S. and R.P. developed the methods; all authors analyzed the data.

## ACKNOWLEDGMENTS

S.C. and A.S. were supported by fellowship from the University Grants Commission (UGC), India. RP thanks Grant No. EMR/2017/001346 of SERB, DST, India for the computational facility. A.M. was supported by the NIH grant GM121971. We thank C. Miles for useful discussions and help with the manuscript, and M. Kwon for attracting our attention to the problem of centrosomal clustering.

## Supporting Material

### Additional examination of the model behavior

We explored the influence of an additional CS-CS force with the length-dependence different from the exponential one: it was shown theoretically that when antiparallel MTs emanating from two CSs overlap, and crosslinking molecular motors at the overlap initiate MT sliding, that the resulting force *f*_*overlap*_ and corresponding potential energy of moving a CS from distance *r*_1_ to *r*_2_ keeping the other CS fixed are :

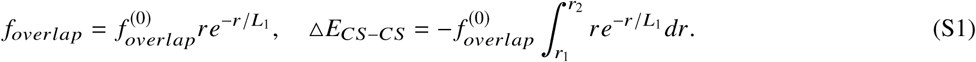

We tested what happens if the CSs interact by a combination of the attractive and repulsive forces. The former can arise when MTs from one CS are reeled in by minus-end directed Dynein motors on another CS. The latter can be generated by plus-end directed Kinesin-5 motors on the antiparallel overlaps between the MTs extending from the interacting CSs. In the first case, the attractive force decreases with distance exponentially, while in the second case, the force is small at small and great distances and is maximal at an intermediate distance. As expected, there has to be a linearly proportional balance between the attractive and repulsive forces for the bipolar spindle to emerge (Fig. S4A,B). If, instead of plus-end directed Kinesin-5 motors, minus-end directed Kinesin-14 motors are dominant at the MT overlaps, then forces generated by both Dynein and Kinesin-14 become attractive, but their distance-dependencies are different. In that case, the motors complement each other: if one is weaker, another one has to be stronger for the bipolar spindle to emerge (Fig. S4D).

The bipolar spindles emerge under many different force combinations, albeit with different frequencies. We tested how the length (pole-to-pole distance) of these spindles depended on the force combinations. Fig. S3A,B demonstrates that to have a proper spindle length (comparable with the cell size), the only condition is that CS-chromosome arm repulsion must be present; other forces do not matter. (They, of course, affect the probability of the bipolarity.) This is one more demonstration of the crucial role of the CS-chromosome arm repulsion.

Chromosome number varies depending on the cell type, and so we explored how the bipolarity depends on the chromosome number. We found that the percentage of the bipolar spindles decreases with the decreasing chromosome number (Fig. S2D). The reason is that smaller numbers of chromosome arms do not repel the CSs strong enough lowering the energy barrier between the mono- and bi-polar configurations. Another reason is that stochastic effects increase with the decreasing number of interacting bodies, effectively enhancing probability of converging to the lowest energy, monopolar configuration.

### Calculation of energy for different spindle configurations

Each of the evolved spindles corresponds to a local minimum in the mechanical energy landscape. We took all mono-, bi- and multi-polar spindles evolved in the simulations with various force combinations, and computed the mechanical energies of each respective configuration of CSs and chromosomes obtained at the end of each stochastic simulations by using the following algorithm. As the chromosomes do not interact with each other, except by the steric repulsion, we start with the chromosomes distributed in the cell exactly as they are at the end of a simulation, with no CSs in the cell, and assign zero energy to this initial condition. Let {*r*_*i*_} be the position of the CSs and {*d*_*ij*_} to be the Euclidean distance between two CSs positioned at *r*_*i*_ and *r*_*j*_ respectively at the end of the simulation. We then bring the CSs one by one from ∞ to *r*_*i*_.

1. The following algorithm is used to evaluate the total energy due to the inter-CS attraction.
  a. We evaluate the potential energy of bringing the *i*^*th*^ CS from ∞ to *r*_*i*_ in presence of other (*i* - 1) CSs which are kept fixed at their individual locations as follows:

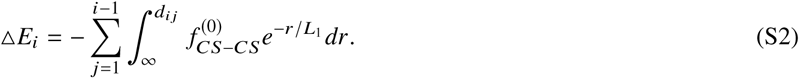
  b. We perform the same task for all CSs and compute the net energy by summing over all the Δ*E*_*i*_s. Note that if Δ*E*_*ij*_ is the energy of interactions of *i*^*th*^ and *j*^*th*^ CSs, this algorithm gives us the sum of all pairwise interaction energies as follows:

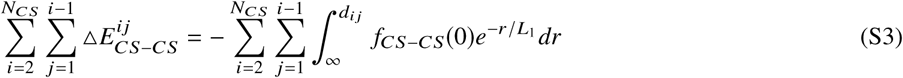
2. We evaluate the total energy of the CS-KT attraction as follows:

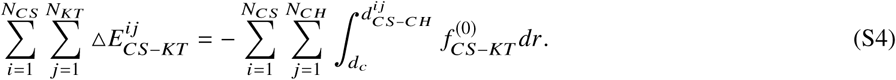 Here, 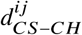 is the distance between the *i*^*th*^ CS and the *j*^*th*^ chromosome. As the CS-KT attraction is constant and acting only inside the cell, the cutoff distance, *d*_c_, for the effective CS-KT attraction is chosen as the diameter of the cell. As we have the elliptical cell with semi-major and semi-minor axes 20, 15, 15 *μ*m, we considered the cutoff distance *d*_*c*_ = 2 × 20 *μ*m= 40 *μ*m in the simulation.
3. We evaluate the total energy of the CS-chromosome arm repulsion as follows:

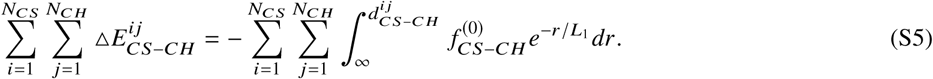
4. We evaluate the total energy of the CS-cortex attraction as follows:

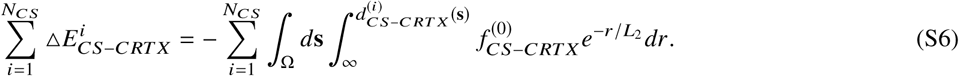 Here, 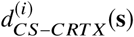 is the distance between *i*^*th*^ CS and the point with coordinate **s** on the cortex.

The average computed energies are reported in Fig. S4G. There are two nontrivial lessons from Fig. S4G: first, the energy differences, under any given force combination, between mono-, bi- and multi-polar spindles are much smaller than the average energy of all spindles under given conditions. This does not mean that the cell can easily change the spindle geometry or that significant stochasticity can destabilize the spindle, because energy barriers between the spindles could be significant. The second lesson is that under all but one condition, the monopolar spindle has the lowest energy (which does not necessarily mean that such spindle is the most stable), followed by the bipolar, and the multipolar spindle. The only case when the bipolar spindle has the lowest energy is when the forces of the CS-CS and CS-cortex attractions are absent.

### Analysis of the energy change upon continuous shift of one CS along the spindle axis and through the ring of chromosomes

We consider the chromosomes to be spatially organized in an annular shaped ring of inner radius *r*_*a*_ and outer radius *r*_*b*_ (Table S1) within a spherical cell of radius *R*_*cell*_ (Table S1). Two CSs are placed on the axis through the center of the ring, perpendicular to the plane into which the ring is embedded into. The chromosome density in the annular ring can be written as 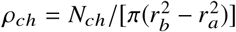. The CS-chromosome arm repulsion can be evaluated as follows:

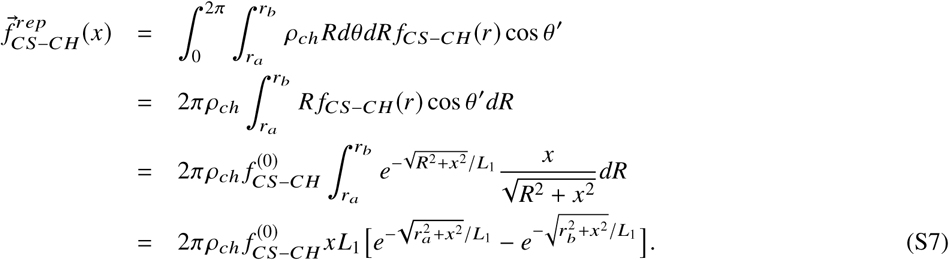

The energy to bring over a CS from ∞ to *x* under the influence of the above mentioned force field is:

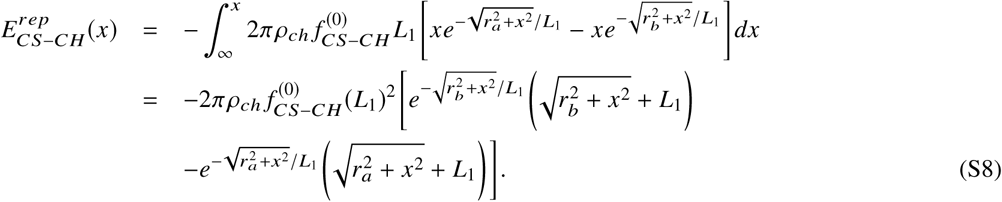

Similarly, we compute the net force on the CS due to the constant CS-KT attraction as follows. KTs are uniformly arranged in a circle of radius R (Table S1), we can define the density of KTs *ρ*_*KT*_ = *N*_*ch*_/*πR*^2^. Then:

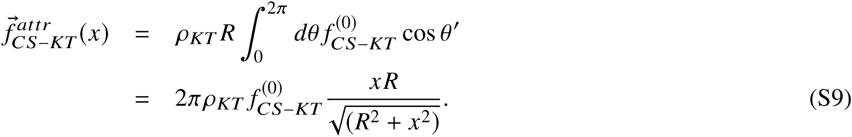

The energy of bringing a CS under the influence of this force field from ∞ to *x* is:

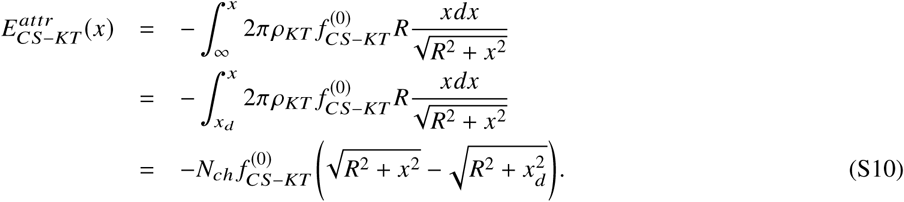

Here we assume that the CS-KT attraction is constant for *x* ≤ *x*_*d*_ (Table S1), elsewhere it vanishes.

If a CS is fixed at *x*′and another CS is placed at *x*, the inter-CS attraction can be written as:

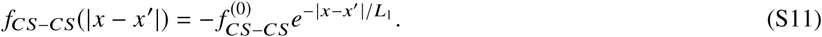

The energy of bringing one CS to *x* from ∞ in the presence of another CS fixed at *x*′ is therefore:

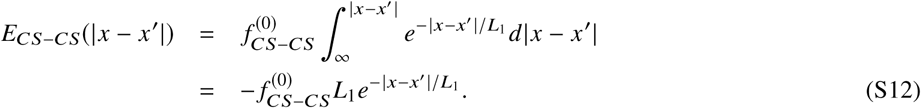

The total energy *E*_*tot*_(*x*) is obtained by summing the energies of CS-chromosome arm repulsion, CS-KT attraction and inter-CS attraction, respectively:

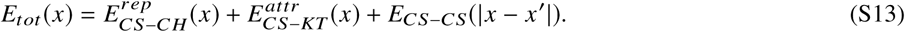

We plot the total energy *E*_*tot*_(*x*) as a function of *x*, the axial distance between the CS and the center of the circular chromosomal ring (Fig. 5G, Fig. S4H).

## SUPPORTING FIGURES

**FIGURE S1.**
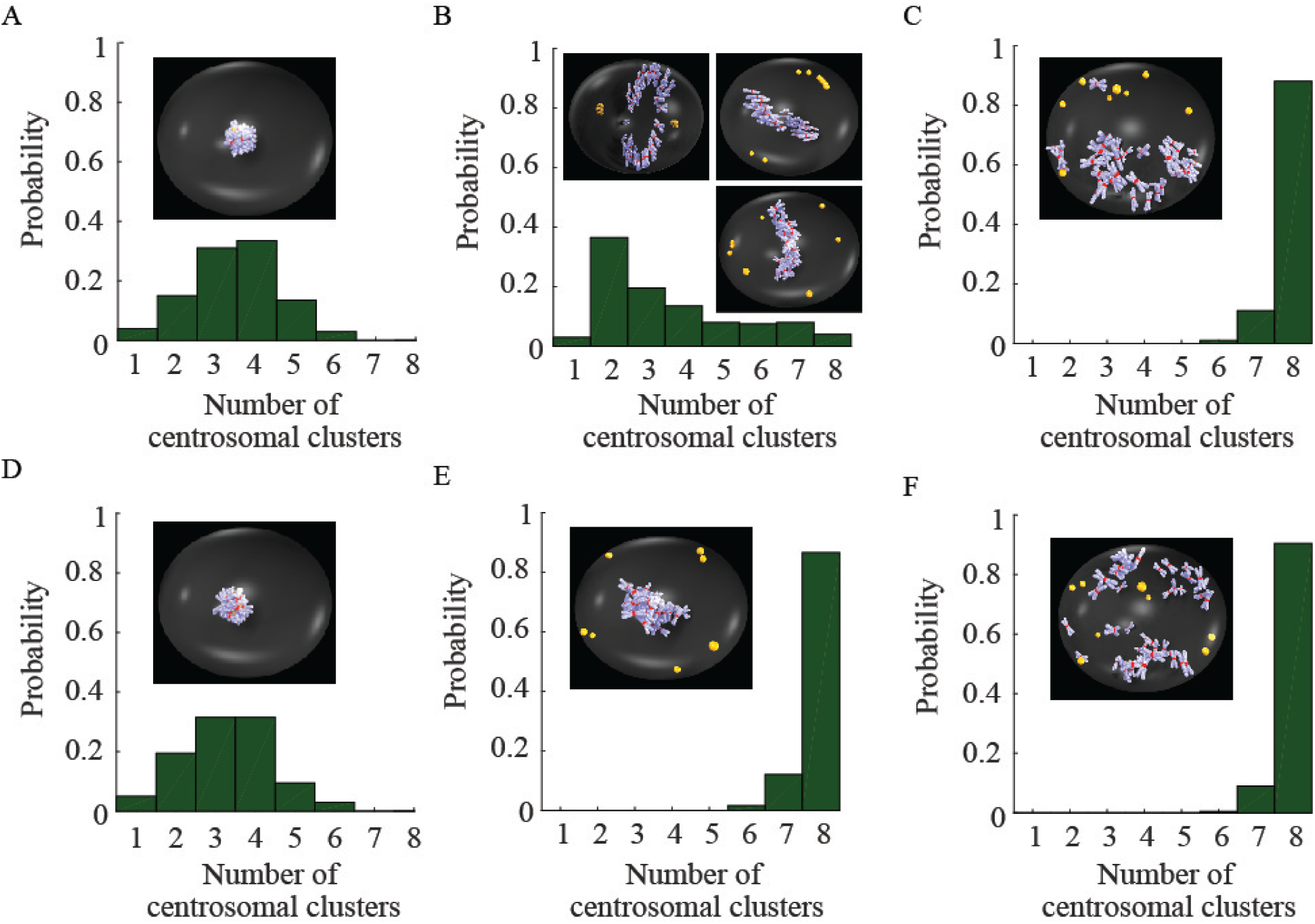
Spindle assembly in the cellular confinement in the absence of the CS-cortex attraction and without CS-CS attraction. CSs are yellow; chromosome arms are blue-and-white; KTs are red. (A) Collapsed spindle in the presence of the CS-KT attraction and in the absence of the inter-CS interaction. (B) Non-robust emergence of the bipolar spindle in the presence of the CS-KT attraction, CS-chromosome arm repulsion and in the absence of the inter-CS interaction. (C) Multipolar spindles develop under the sole influence of CS-chromosome arm repulsion. (D) Collapsed spindle when the CS-KT attraction is supplemented by the inter-CS repulsion. (E) Multipolar spindles develop when the CS-KT attraction and CS-chromosome arm repulsion are supplemented by the inter-CS repulsion. (F) Multipolar spindles develop when the CS-chromosome arm repulsion is combined with inter-CS repulsion.

**FIGURE S2.**
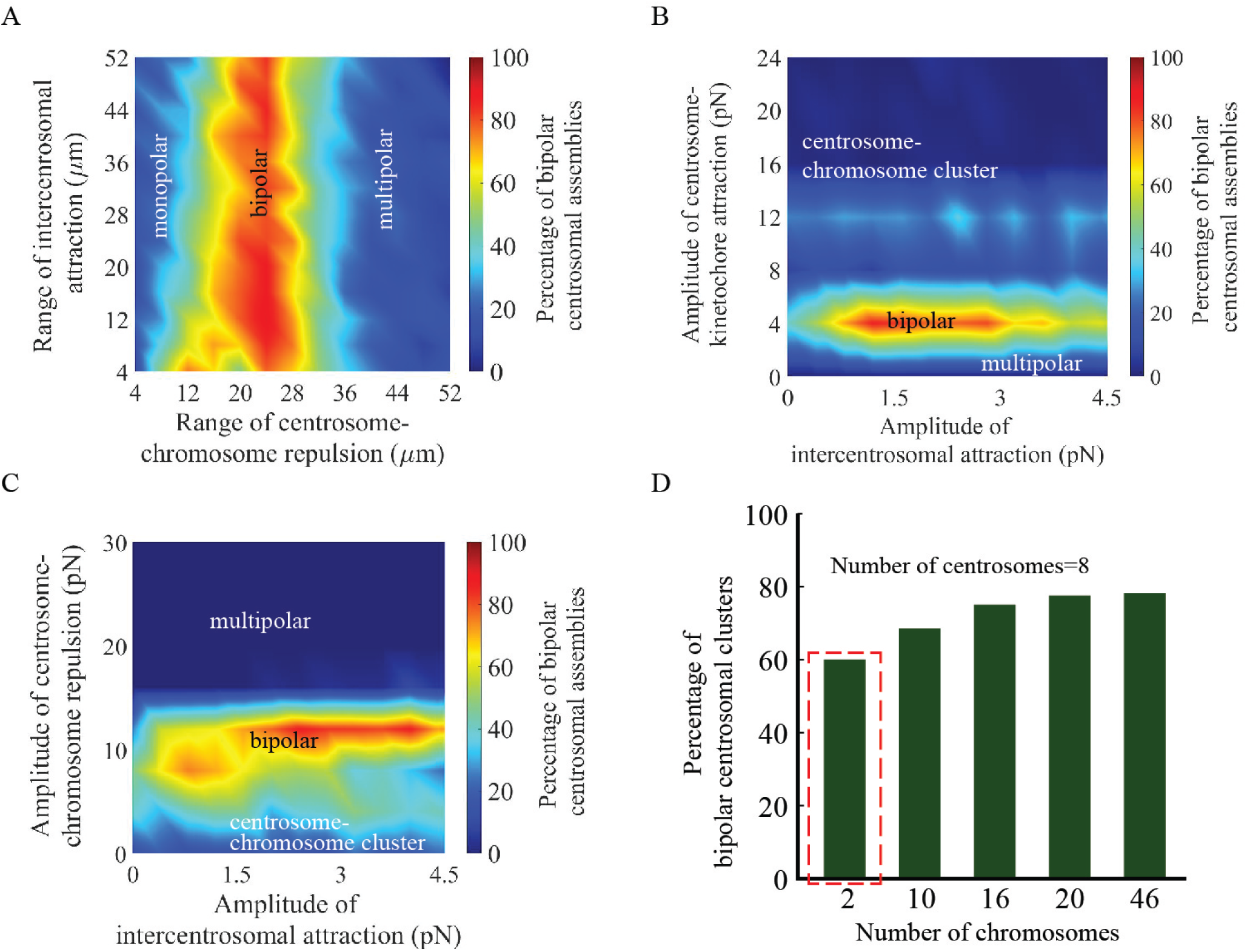
Model sensitivity to the parameters. (A) The bipolarity is sensitive to variations of the range of CS-chromosome arm repulsion and insensitive to that of the CS-CS attraction. (B) The bipolarity is sensitive to the amplitude of the CS-KT attraction but insensitive to that of the CS-CS attraction. (C) The bipolarity is sensitive to the amplitude of the CS-chromosome arm repulsion but insensitive to that of the CS-CS attraction. (D) Percentage of bipolar spindles depending on the number of chromosomes. The statistics for two chromosomes (enclosed by red dashed rectangle) is obtained by changing the amplitudes of the CS-CS attraction and of the CS-cortex attraction to 1.6 pN and 0.36 pN respectively from the base parameters.

**FIGURE S3.**
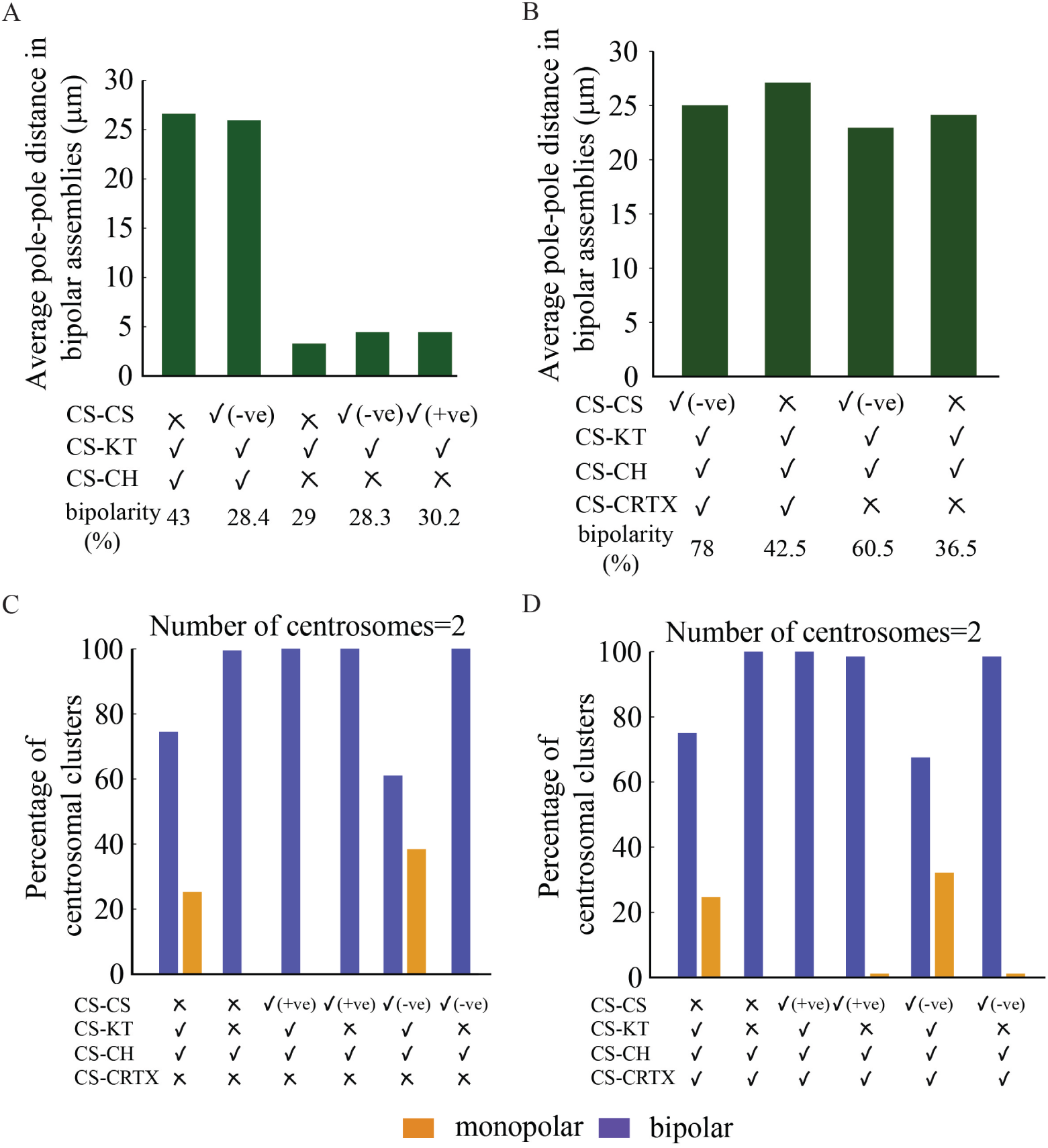
Dependence of the spindle characteristics on the force combinations. (A) Average pole-pole distance between two CS clusters in the bipolar spindles assembling under the influence of various force balances in unconfined geometry. **✓**(**✗**) indicates presence (absence) of a particular interaction. -ve (+ve) denotes attractive(repulsive) forces. (B) between two CS clusters in the bipolar spindles assembling under the influence of various force balances in confined geometry. (C-D) The statistics of the mono/bi-polar spindles for various combinations of forces in confined geometry in the absence (C) or presence (D) of the centrosome-cortex attraction.

**FIGURE S4.**
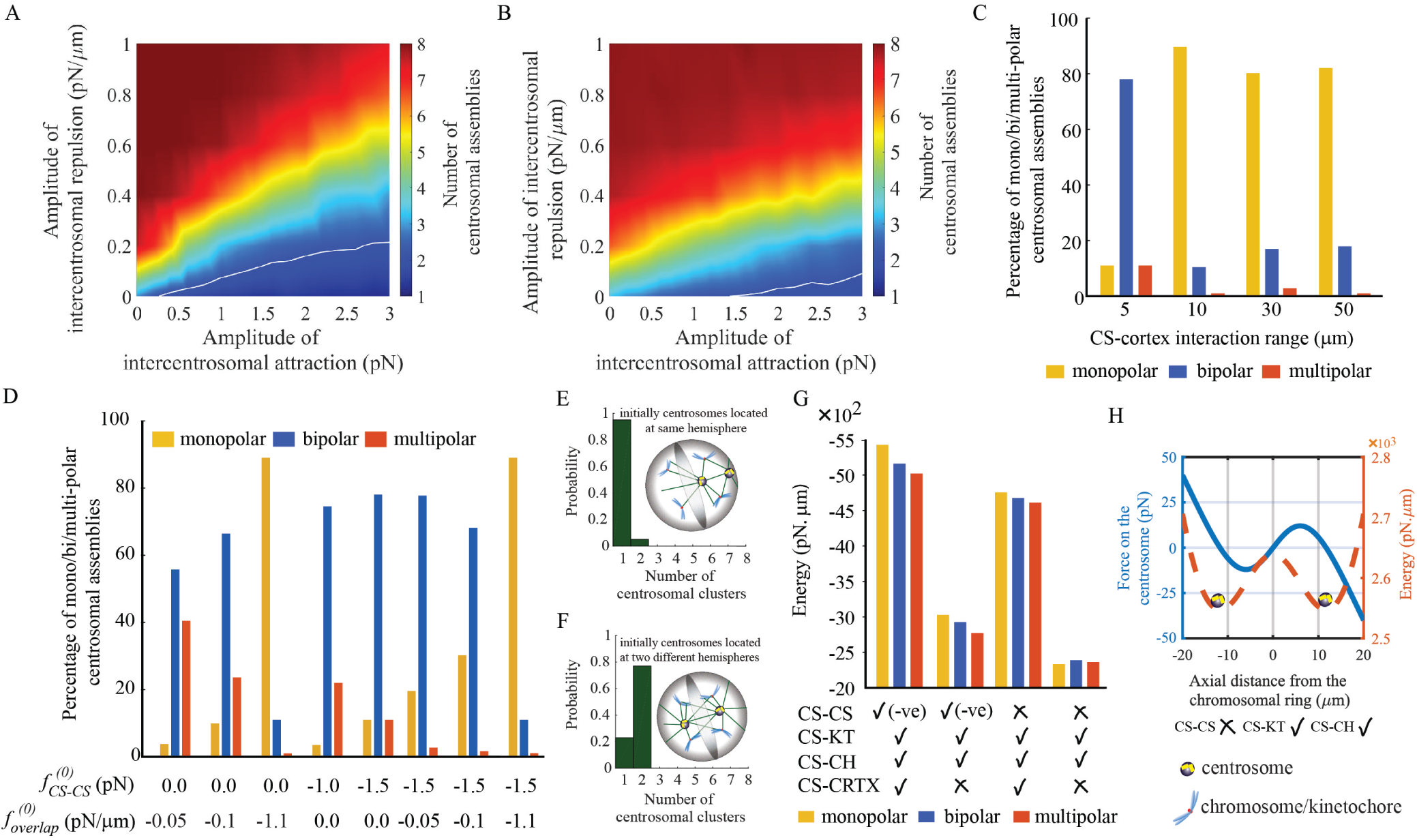
Additional spindle characteristics. (A-B) Bipolar centrosomal assembly contour (white solid line) shows the optimized balance of the inter-CS repulsion and attraction in unconfined (A) and confined (B) geometry. (C) Percentage of mono/bi/multi-polar spindles depending on the interaction range *L*_2_. (D) Percentage of mono/bi/multi-polar spindles depending on relative strengths of ‘end-on’ (*f*_*CS*−*CS*_, dynein) and ‘overlap’ (*f*_*overlap*_, kinesin-14) attractive inter-CS forces. (E-F) Percentage of mono/bi-polar spindles in 2-CS cells depending on whether two CSs are initially localized in the same (E) or different (F) hemispheres. The shaded equatorial plane distinguishes the hemispheres as shown in the inset. (G) Average spindle energies depending on the combinations of various forces in cellular confinement. (H) Dependence of the force and energy on the CS-CS distance in the thought experiment corresponding to Fig. 5G balance landscape in the absence of the inter-CS attraction.

## SUPPORTING MOVIES

**M1**. Dynamic exploration of the energy space in the presence of intercentrosomal attraction, centrosome-kinetochore attraction and centrosome-chromosome arm repulsion in unconfined geometry results in the monopolar spindle. CSs are yellow; chromosome arms are blue-and-white; KTs are red.

**M2**. Dynamic exploration of the energy space in the presence of intercentrosomal repulsion, centrosome-kinetochore attraction, centrosome-chromosome arm repulsion and centrosome-cortex attraction results in the multipolar spindle. CSs are yellow; chromosome arms are blue-and-white; KTs are red.

**M3**. Dynamic exploration of the energy space in the presence of intercentrosomal attraction, centrosome-kinetochore attraction, centrosome-chromosome arm repulsion and centrosome-cortex attraction results in the bipolar spindle. CSs are yellow; chromosome arms are blue-and-white; KTs are red.

**Table S1.** List of model parameters.

| Parameters | Description | Value |
| --- | --- | --- |
| $f_{CS-CS}^{(0)}$ | amplitude of intercentrosomal attraction (repulsion) | -1.5 (1.5) pN |
| $f_{overlap}^{(0)}$ | amplitude of intercentrosomal attraction (repulsion) | $\sim \pm 1$ pN/μm |
| $f_{CS-KT}^{(0)}$ | amplitude of centrosome-kinetochore attraction | -4.0 pN |
| $f_{CS-CH}^{(0)}$ | amplitude of centrosome-chromosome arm repulsion | 10 pN |
| $f_{CS-CRTX}^{(0)}$ | amplitude of the centrosome-cortex attraction | -0.4 pN/μm<sup>2</sup> |
| $L_1$ | range of CS-CH and CS-CS interactions | 20 μm |
| $L_2$ | range of CS-cortex interaction | 5 μm |
| $L_{merge}$ | merging distance below which centrosomes are considered clustered | 1.5 μm |
| $\beta$ | inverse temperature | 100 (pN × μm)<sup>-1</sup> |
| additional parameters for analytical model |  |  |
| $r_a, r_b$ | inner and outer radius of the annular chromosome arrangement | 12, 16 μm |
| $R$ | radius of the circular ring where kinetochores are arranged | 14 μm |
| $R_{cell}$ | radius of the cell | 20 μm |
| $x_d$ | cut off distance for kinetochore fiber attraction | 20 μm |

**TABLE S2.**
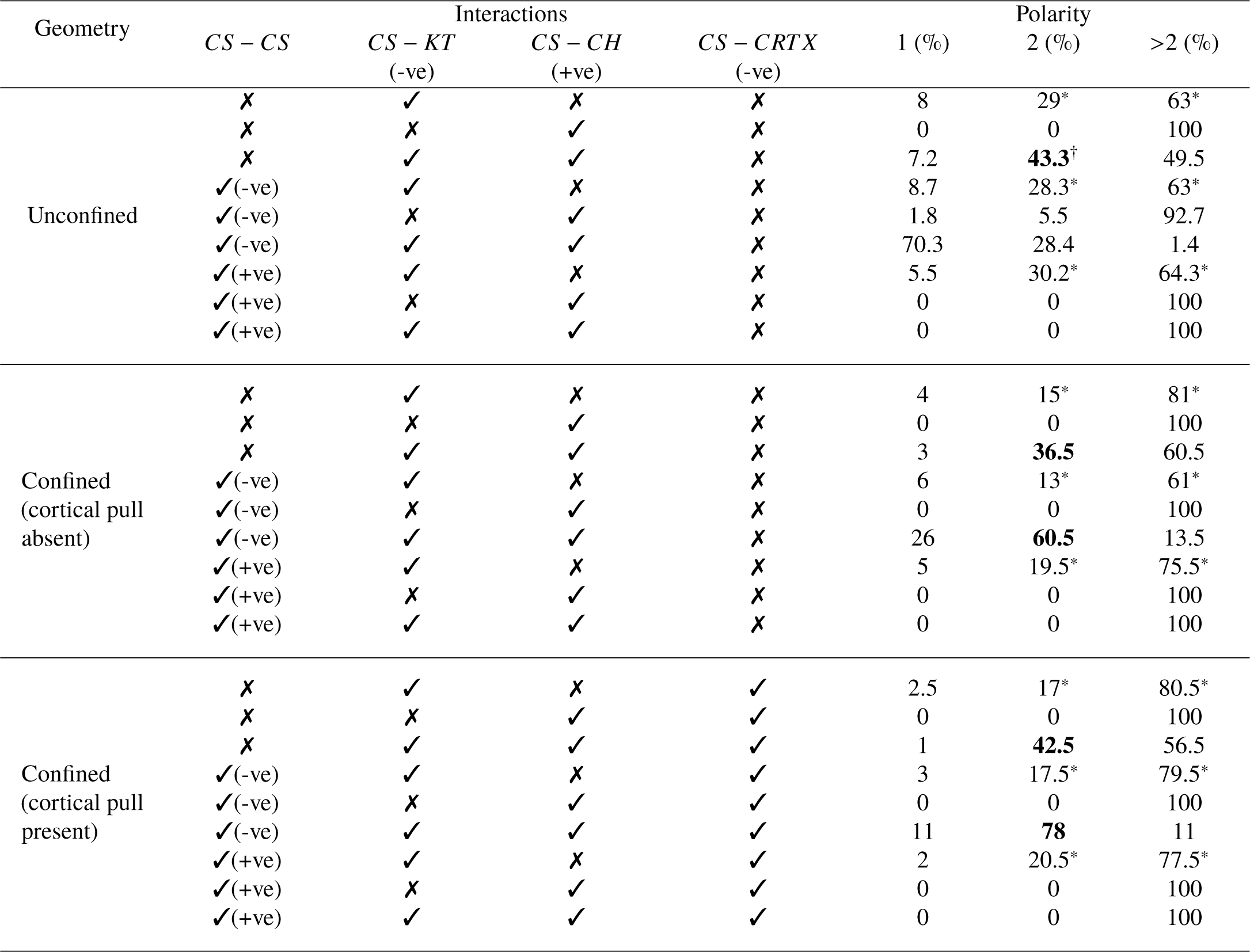
Statistics of the CS clusters. ‘-ve’ (‘+ve’) denotes attractive (repulsive) force fields. * Nearly collapsed CS-chromosome aggregate with 2 or more CS clusters. ^†^ A mixture of proper bipolar configurations with chromosomes at the midzone between two CS clusters and abnormal structures with 2 CS clusters.

